# Modeling in yeast how rDNA introns slow growth and increase desiccation tolerance in lichens

**DOI:** 10.1101/2021.05.01.442275

**Authors:** Daniele Armaleo, Lilly Chiou

**Affiliations:** Department of Biology, Duke University, Durham, North Carolina 27708, USA; Curriculum in Genetics and Molecular Biology, University of North Carolina at Chapel Hill, Chapel Hill, North Carolina 27599, USA

**Keywords:** Desiccation tolerance, rDNA introns, rRNA processing, Ribosome assembly, Lichen fungi, *Cladonia grayi*, *Saccharomyces cerevisiae*, Trehalose, CRISPR

## Abstract

We define a molecular connection between ribosome biogenesis and desiccation tolerance in lichens, widespread symbioses between specialized fungi (mycobionts) and unicellular phototrophs. Our experiments test whether the introns present in the nuclear ribosomal DNA of lichen mycobionts contribute to their anhydrobiosis. Self-splicing introns are found in the rDNA of several eukaryotic microorganisms, but most introns populating lichen rDNA are unable to self-splice, being either degenerate group I introns lacking the sequences needed for catalysis, or spliceosomal introns ectopically present in rDNA. Although all introns are eventually removed from rRNA by the splicing machinery of the mycobiont, Northern analysis of its RNA indicates that they are not removed quickly during rRNA transcription but are still present in early post-transcriptional processing and ribosome assembly stages, suggesting that delayed splicing interferes with ribosome assembly. To study the phenotypic repercussions of lichen introns in a model system, we used CRISPR to introduce a spliceosomal intron from the rDNA of the lichen fungus *Cladonia grayi* into all nuclear rDNA copies of the yeast *Saccharomyces cerevisiae*, which lacks rDNA introns. Three intron-bearing yeast mutants were constructed with the intron inserted either in the 18S rRNA genes, the 25S rRNA genes, or in both. The mutants removed the introns correctly but had half the rDNA genes of the wildtype strain, grew 4.4 to 6 times slower, and were 40 to 1700 times more desiccation tolerant depending on intron position and number. Intracellular trehalose, a disaccharide implicated in desiccation tolerance, was detected, but at low concentration. Overall, our data suggest that the constitutive interference of the intron splicing machinery with ribosome assembly and the consequent lowering of the cytoplasmic concentration of ribosomes and proteins are the primary causes of slow growth and increased desiccation tolerance in the yeast mutants. The relevance of these findings for slow growth and desiccation tolerance in lichens is discussed.

## Introduction

In the study of desiccation tolerance, special attention has been devoted recently to anhydrobiotes, organisms that can survive losing more than 99% of their water (Leprince and Buitink 2015; Koshland and Tapia 2019). We focus here on ribosome biogenesis as a key node in the control of desiccation tolerance in lichens, stable anhydrobiotic symbioses between specialized fungi (mycobionts) and unicellular green algae or cyanobacteria (photobionts). Transcriptomic analyses addressing lichen desiccation tolerance suggest ribosomal involvement (Junttila *et al.* 2013; Wang *et al.* 2015). Several ribosomal assembly and translation functions appear to be under purifying selection in the lichen *Cladonia grayi* (Armaleo *et al.* 2019) suggesting that they play pivotal roles in the anhydrobiotic lichen symbiosis. In yeast, ribosomal network regulation is central to the Environmental Stress Response (ESR) (Gasch *et al.* 2000) and mutations hindering ribosomal assembly enhance desiccation tolerance (Welch *et al.* 2013).

Lichen anatomy is complex for microorganisms (Honegger 2012) but lacks the water management systems found in plants, such as roots, vascular tissues or waxy cuticles. Lichens employ water-storing polysaccharides and evaporation barriers like cortices, hydrophobins, and secondary compounds to slow down water loss but eventually, within minutes or hours of drying depending on conditions, residual metabolism and transcription come to a complete halt throughout their thalli (Junttila *et al.* 2013; Wang *et al.* 2015; Candotto Carniel *et al.* 2020). Yet lichens can survive losing more than 95% of their water content and remaining dehydrated for long periods of time, a stress that would kill most organisms (Kranner *et al.* 2008). This is in contrast with non-lichenized fungi, which spend most of their life cycles protected within their substrates and survive desiccation through spores. This work focuses exclusively on mycobionts. For a review of lichen desiccation tolerance which includes photobionts, see(Gasulla *et al.* 2021).

Frequent desiccation and rehydration induce protein denaturation and aggregation as well as the formation of reactive oxygen species (ROS) that can cause direct damage to DNA, proteins, and lipids. Antioxidants and ROS-processing enzymes, common defenses against stress, are thought to protect also lichens from desiccation damage (Kranner *et al.* 2008). However, the ability of lichens to repeatedly withstand extreme desiccation suggests the involvement of additional defense mechanisms. Thus, we hypothesized (Armaleo *et al.* 2019) that the many rDNA introns present in lichen fungi (DePriest 1993; Gargas *et al.* 1995; Bhattacharya *et al.* 2000; Bhattacharya *et al.* 2002) may provide such an extra defense through their effects on ribosome biogenesis. The introns relevant to this work are spliceosomal introns and group I introns. Spliceosomal introns, universal among eukaryotes, are normally associated with Polymerase II transcription of protein-coding genes, require specialized host factors to be excised, and perform important regulatory roles (Poverennaya and Roytberg 2020). Group I introns are found mostly in the ribosomal DNA of a number of eukaryotic microorganisms, and normally form ribozymes able to fold into a conserved structure that catalyzes self-splicing from the transcript without the aid of host factors *in vitro* (Cech 1990), although self-splicing can be aided by maturase proteins facilitating proper folding of the RNA *in vivo* and *in vitro* (Lambowitz et al. 1999). Experiments with the self-splicing group I intron from the ciliate *Tetrahymena thermophila* in its original host or transferred into yeast rDNA have shown that the intron splices itself co-transcriptionally within seconds (Brehm and Cech 1983), and that 99% of the introns are removed before the end of 35S pre-rRNA transcription (Jackson *et al.* 2006), which takes ~2 minutes per molecule in yeast (Osheim *et al.* 2004). Intron presence has no discernible consequences on growth rate (Lin and Vogt 1998) or other phenotypes (Nielsen and Engberg 1985). This is why group I introns are commonly considered harmless genetic parasites, including the relatively few group I introns found in the rDNA of non-lichen fungi (Haugen *et al.* 2004b; Hedberg and Johansen 2013). Compared to non-lichen fungi, the rDNA of lichen mycobionts contains many group I introns (Fig.1) and a few ectopic spliceosomal introns (Bhattacharya *et al.* 2000) which normally operate only in mRNA.

**Figure 1.**
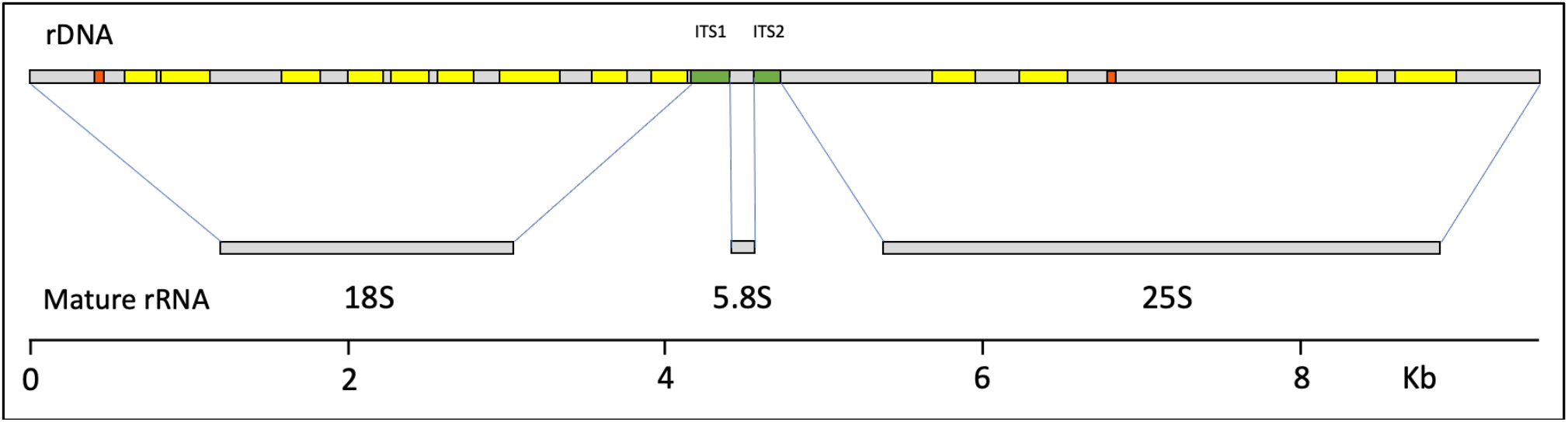
Introns in the rDNA of a single-spore isolate from the lichen fungus *Cladonia grayi* (Armaleo et al. 2019). Top: the 18S, 5.8S, and 25S region of one rDNA repeat; gray, rRNA coding sequences; green, internal transcribed spacers; yellow, group I introns; orange, spliceosomal introns. Middle: the three mature rRNAs produced from this region. Bottom: scale in kilobases. Sequences were retrieved from https://mycocosm.jgi.doe.gov/Clagr3/Clagr3.info.html.

Lichen group I introns tested *in vitro* were found to be “degenerate”, *i.e.* unable to self-splice (DePriest and Been 1992; Haugen *et al.* 2004b). This lack of self-splicing has remained a puzzling observation which we reframe here as a central feature of lichen rDNA introns and a fundamental key to interpret our yeast results and extrapolate them to lichens. To this end, Figure 2 combines self-splicing data across several lichen and non-lichen fungi with their group I intron lengths and includes also group I introns lengths from other microbial eukaryotes. Lichen group I introns, in which all self-splicing tests were negative, are the shortest and lack an average of 100 nucleotides relative to group I introns from non-lichen fungi. The lost sequences have been shown to be necessary for catalysis (Haugen *et al.* 2004b). Since such degenerate introns are completely removed from mature rRNA (Fig. 1), a specific group I intron splicing machinery must have evolved in lichens. Since spliceosomal introns require spliceosomes for removal, the need for a dedicated splicing machinery is a characteristic shared in lichens by group I and spliceosomal rDNA introns. There are occasional self-splicing group I introns in lichen fungi (Haugen *et al.* 2004a; Reeb *et al.* 2007), which would quickly splice themselves out of any significant interference with rRNA processing. Those rare cases do not negate that loss of self-splicing is a prevalent feature of lichen rDNA introns.

**Figure 2.**
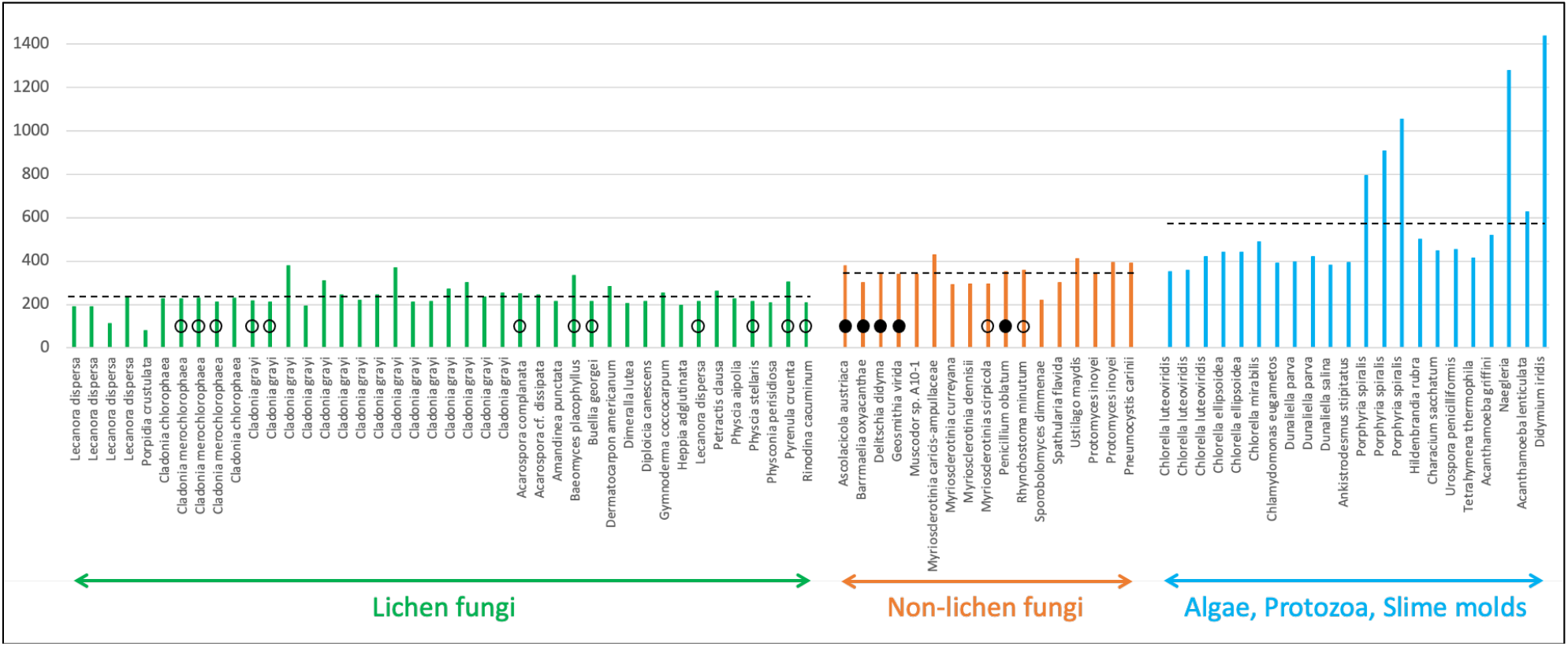
Length and splicing comparisons between group I introns from lichen and non-lichen fungi. Each colored vertical bar represents the length of an intron (nucleotide number on the ordinate). Dotted lines represent the average intron lengths within each of the three groupings (233, 339, and 586 nucleotides respectively). Introns tested for self-splicing *in vitro* by (DePriest and Been 1992) and (Haugen *et al.* 2004b) are highlighted with open circles (no splicing) or filled circles (splicing). Length data were compiled from (Gargas *et al.* 1995), (Haugen *et al.* 2004b), (Armaleo *et al.* 2019). This is not a comprehensive compilation of all available group I intron data.

The hypothesis prompting us to test the effect of lichen introns on desiccation tolerance is that the splicing machinery necessary to remove them from nascent rRNA interferes with rRNA processing and ribosome assembly (Fig. 3), leading to slow growth and to increased desiccation tolerance. A Northern analysis of intron splicing in the cultured lichen mycobiont *Cladonia grayi* supports that hypothesis. To test in a model system the effects of non-self-splicing introns on growth and desiccation tolerance, we used CRISPR to transfer a spliceosomal rDNA intron (first on left in Figure 1) from the lichen *Cladonia grayi* into all 150 rDNA repeats of *S. cerevisiae* (Chiou and Armaleo 2018), which lacks rDNA introns. We inserted the intron at two yeast rDNA sites occupied by introns in *C. grayi* (Figure 3 shows the intron inserted at the 18S site). Three intron-bearing mutants were constructed, with the intron stably inserted either in all copies of the 18S rRNA gene, of the 25S rRNA gene, or of both. Among wild type and mutant strains, we compared ribosomal DNA repeat number, growth rates, desiccation tolerance, cell morphology and intracellular concentration of trehalose, a disaccharide associated with desiccation resistance in yeast and other organisms (Koshland and Tapia 2019). The main effects of the introduced introns were dramatic decreases in growth rate and equally significant increases in desiccation resistance.

**Figure 3.**
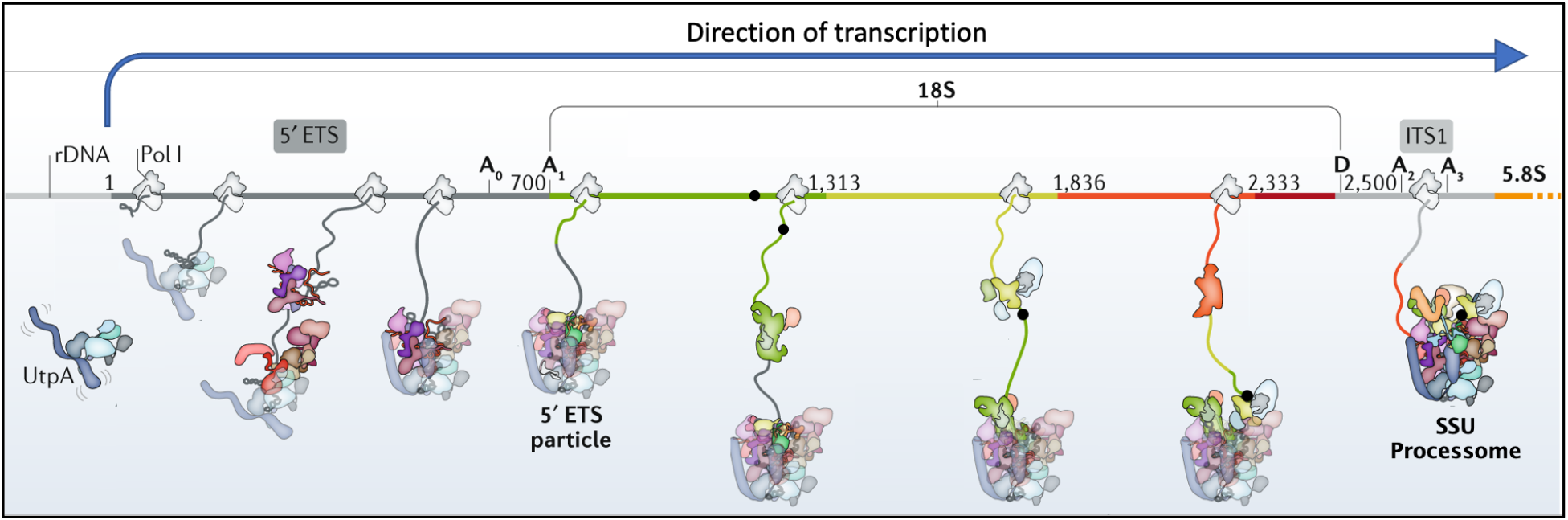
Visualization of how delayed intron splicing could interfere with ribosome assembly. The scheme depicts early nucleolar phases of the cotranscriptional assembly of yeast 18S pre-ribosomal particles. From the start of transcription, several assembly factors interact dynamically with the nascent rRNA, initiating ribosome construction. The grey parts of the nascent RNA correspond to the 5’ ETS and ITS1, the colored parts to the mature 18S rRNA. The spliceosomal intron we inserted into the yeast 18S rDNA is depicted as a proportionally sized black dot located on the DNA and incorporated upon transcription into the growing pre-ribosomal particle. The bulky and relatively slow spliceosome (not depicted here, but comparable in size to the processome) removing the intron is expected to interfere with ribosome assembly, delaying it. We depicted the intron as persisting up to the processome stage (as inferred from a Northern blot involving a different 18S intron; see Results). A similar interference by the intron is expected to delay maturation of 25S pre-ribosomal particles (not shown). The diagram was modified from (Klinge and Woolford 2019) with permission.

## Materials and Methods

### Plasmids, strains, and media

The lichen mycobiont used for the Northern is the *C. grayi* single-spore isolate Cgr/DA2myc/ss (Armaleo *et al.* 2019), routinely propagated on MY medium (Hamada 1996). DH5α *E. coli* cells containing the plasmid pCAS were obtained from Addgene (#60847). Plasmid pCAS (8.7 Kb) is a kanamycin/G418 shuttle vector carrying the gene for Cas9 and a generic guide RNA expression cassette (Ryan and Cate 2014). Competent *E. coli* strain DH10B (New England Biolabs, C3019I) was used for transformations. *E. coli* was grown in LB medium (Protocols 2006) at 37°C. For *E. coli* transformant selection, kanamycin (ThermoFisher Scientific) was added to the LB medium to 100 mg/L. The yeast strain used was *S. cerevisiae* YJ0 (*MAT**a**, gal4*Δ, *gal80*Δ, *ura3–52, leu2–3, 112 his3, trp1, ade2–101*) (Stafford and Morse 1998), and was typically grown in YEPD medium (Protocols 2010) at 30°C. To select for yeast containing the transforming plasmid, G418 (VWR) was added to the plate medium to 200 mg/L.

### Northern of group I intron splicing in the lichen fungus

To prepare total genomic RNA, small volumes of a *C. grayi* mycobiont liquid culture were seeded onto 47-mm diameter, 20 μm pore size Nylon filters. Filters were placed on MY plates and the mycobiont was grown for 1 month at room temperature. Dry weight of mycelia per filter was determined separately, two fresh filters (total dry weight equivalent ~ 10 mg) were removed from plates, wetted in RNAlater (Ambion), mycelia were scraped off, pooled, and total genomic RNA was extracted as described in (Armaleo *et al.* 2019). Using standard Northern protocols, 1% agarose gel electrophoresis was performed with 1 μg of total RNA per lane, and the RNA was then transferred and UV-crosslinked to a Hybond-N+ nylon membrane (Amersham). To prepare the probe, primers CgInt3F and CgInt3R (Table 1) were used to isolate by PCR (Table 2A) from *C. grayi* DNA a 109 bp fragment internal to group I intron-3 (third from left in Figure 1). The PCR fragment was cloned into the pGEM-T Easy vector (Promega), whose polycloning site is flanked by two phage promoters (T7 and SP6) for RNA probe synthesis. Sequence and orientation of the insert were confirmed by sequencing. The plasmid was linearized with SacII, and SP6 polymerase was used for transcription and digoxigenin (DIG) for labeling of the intron-3 RNA antisense probe, using the Roche DIG Northern Starter Kit. The same kit was used for hybridization and signal detection on X-ray film.

**Table 1.**
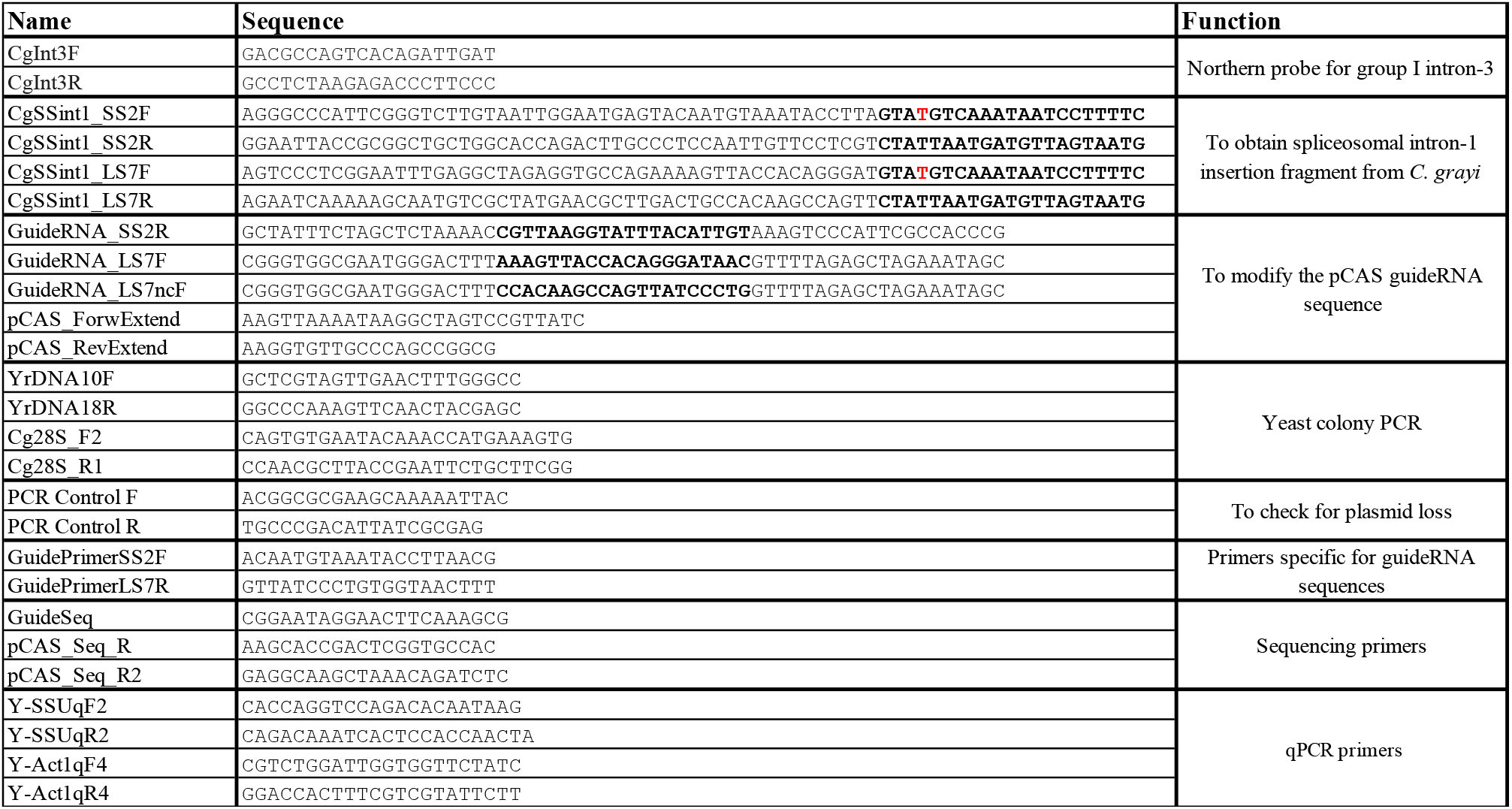
Primers used in this study. The 3’ end sequences bolded in the first set of primers anneal to the 5’ and 3’ end regions of the intron respectively. The T in red is a mismatch designed to modify the 5’ splice site. The fifty 5’ bases match the yeast rDNA flanking the Cas9 cut site. The twenty bases bolded in the second set of primers are the guide sequences for Cas9.

**Table 2.** Thermocycling conditions referred to in the text.

| A |  | B |  | C |  | D |  | E |  | F |  |
| --- | --- | --- | --- | --- | --- | --- | --- | --- | --- | --- | --- |
| Temp. | Time | Temp. | Time | Temp. | Time | Temp. | Time | Temp. | Time | Temp. | Time |
| 94°C | 4 min | 98°C | 30 sec | 98°C | 30 sec | 94°C | 4 min | 95°C | 5 min | 95°C | 60 sec |
| *94°C | 30 sec | *98°C | 10 sec | *98°C | 10 sec | *94°C | 30 sec | *95°C | 30 sec | *95°C | 30 sec |
| 55°C | 30 sec | 58°C | 1 min | 55°C | 30 sec | 50°C | 30 sec | 60°C | 30 sec | 56°C | 30 sec |
| 72°C | 30 sec | 72°C | 4 min | 72°C | 10 sec | 72°C | 15 sec | to * 39x |  | to * 39x |  |
| to * 39x |  | to * 30x |  | to * 34x |  | to * 39x |  | 72°C | 2 min |  |  |
| 72°C | 7 min | 72°C | 2 min | 72°C | 5 min | 72°C | 7 min |  |  |  |  |

### Guide sequence introduction into pCAS by inverse PCR, screening, plasmid isolation

In our hands the yields were very low when we used the procedure recommended by (Ryan *et al.* 2016) to introduce a desired guide sequence into pCAS. Their procedure involves PCR with two self-complementary “guide RNA” primers, followed by DpnI restriction to eliminate the original methylated plasmid before transformation. We therefore decided to update the inverse PCR procedure developed for plasmid mutagenesis by (Hemsley *et al.* 1989). We found that by using two primers which are not self-complementary dramatically increases product yield and eliminates the need for DpnI treatment (Fig. 4). One of the two is 60 bp long and is the mutagenic primer (GuideRNA primers in Table 1), designed according to the guidelines in (Ryan *et al.* 2016) to contain the desired 20-bp guide sequence flanked on both sides by 20-bp sequences homologous to the plasmid. The other is a normal, shorter PCR primer not overlapping with but immediately adjacent to the long primer on the plasmid sequence (Fig. 4). Using the inverse PCR method, we constructed three modified pCAS plasmids, pCAS_SS2, pCAS_LS7, and pCAS_LS7nc with, respectively, primer pairs GuideRNA_SS2R and pCAS_ForwExtend, GuideRNA_LS7F and pCAS_RevExtend, and GuideRNA_LS7nc and pCAS_RevExtend (Table 1). For high fidelity PCR, we used Phusion HF DNA Polymerase (NEB) according to the manufacturer’s specifications. Each 10-μl PCR contained 1 ng pCAS plasmid. It is important to use the relatively high dNTP concentration in the reaction (200 μM each) recommended by the manufacturer to avoid unwanted deletions and mutations around the plasmid ligation junction. At lower dNTP concentrations the 3’>5’ exonuclease activity of Phusion polymerase increases and could damage primer and amplicon ends. Thermocycling conditions were as described in Table 2B. The linear PCR product (Fig. 4) was 5’ phosphorylated and blunt-end-ligated using standard protocols. Competent *E. coli* (strain DH10B) were transformed with each of the three plasmids and plated on LB-kanamycin medium. Colony PCR was first used to screen 10-20 transformants for the presence of the correct guide sequence. The primers used for colony PCR (Table 1) were GuidePrimerSS2F, and pCAS_Seq_R for plasmid pCAS-SS2, GuidePrimerLS7R and GuideSeq for plasmid pCAS-LS7, and GuideSeq and pCAS_Seq_R2 for plasmid pCAS-LS7nc. For the first two plasmids, one primer (GuidePrimerSS2F or GuidePrimerLS7R) was specific for the guide sequence so that a PCR product would form only in presence of the correct guide sequence. Both primers used for plasmid pCAS-LS7nc were flanking the guide sequence, whose correctness was then directly confirmed by sequencing. PCR conditions were as described in Table 2D. Plasmids were isolated from transformants, and presence of the correct gRNA sequence was confirmed by sequencing using forward primer GuideSeq and either pCAS_Seq_R or pCAS_Seq_R2 as the reverse primer.

**Figure 4.**
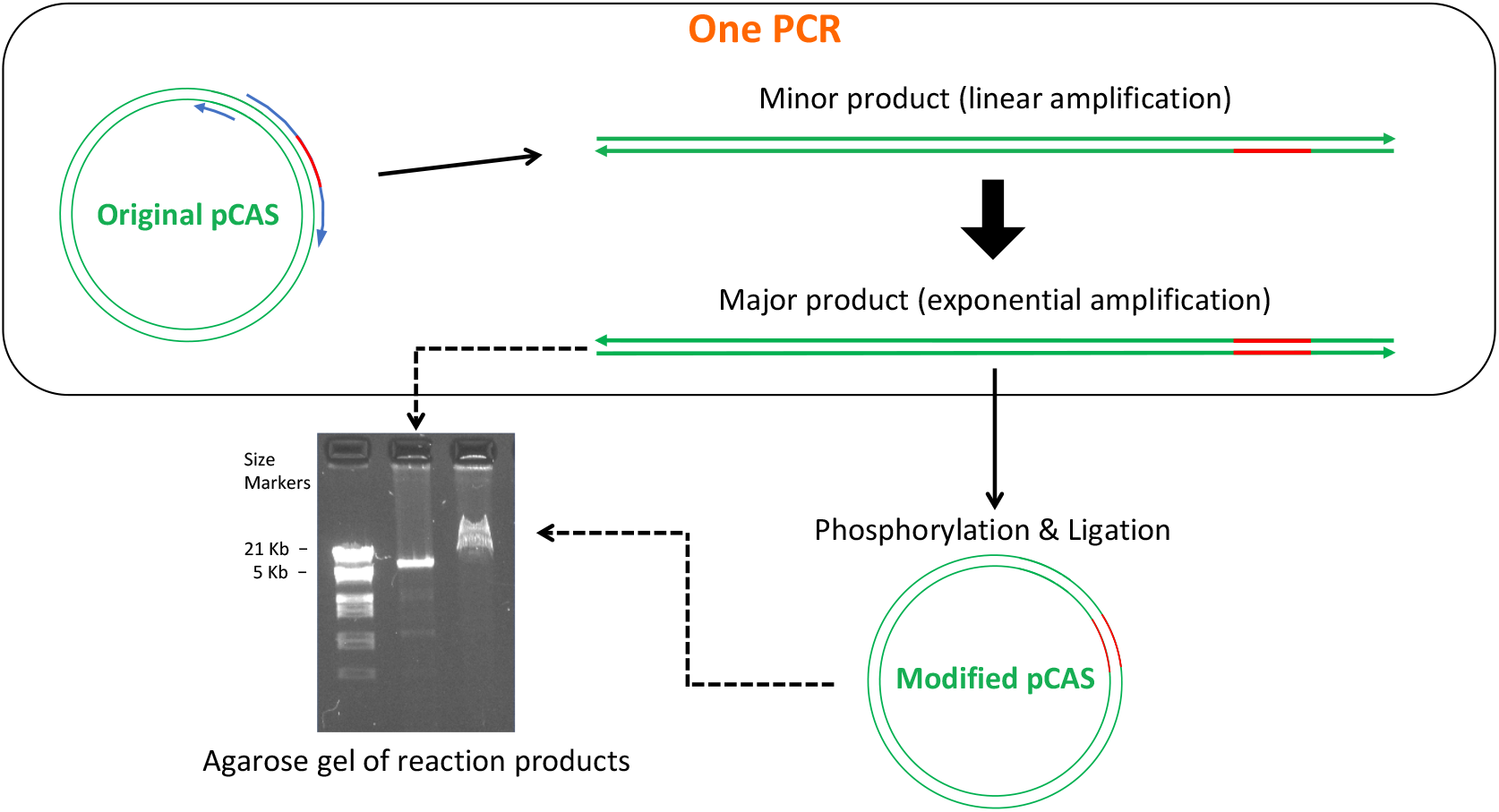
Inverse PCR is fast and efficient in modifying CRISPR guide sequences. Drawings are not to scale and primer sizes (blue rounded arrows on original pCAS) are exaggerated relative to the size of the plasmid (green double lines). The modified guide sequence is colored orange on primers and plasmids. Components and events in the single PCR reaction are enclosed within the rounded rectangle at the top. Either one of the two possible primer arrangements on the original pCAS can be chosen to create the same modification of the guide RNA. The figure depicts only one primer arrangement. The 5’ nucleotide of the 60 base-long GuideRNA primer and of the 20 or 27 base-long Extend primer (Table 1) correspond to adjacent nucleotides on the plasmid sequence, producing full-length, blunt-ended, linear products that are directly phosphorylated, ligated, and used in transformation. Sub-nanogram quantities of the original 8.7 Kb plasmid can yield micrograms of modified plasmid as shown by the gel image. The very large excess of modified plasmid over the amounts of original and minor product DNA bypasses the need to eliminate the minor DNAs before transformation, as 50% or more of the *E. coli* transformants will have the desired guide sequence.

### Intron PCR

*Cladonia grayi* mycobiont DNA was isolated as described in (Armaleo and May 2009). The 57-bp intron (first on left in Figure 1) we chose for transfer into yeast rDNA was amplified (Phusion HF DNA Polymerase) from 5 ng *C. grayi* DNA with two primer pairs, CgSSint1_SS2F and CgSSint1_SS2R for insertion into the 18S (SSU) sequence, or CgSSint1_LS7F and CgSSint1_LS7R for insertion into the 25S (LSU) sequence (Fig. 5 and Table 1). PCR conditions are listed in Table 2C. PCR products were cleaned with a QIAquick PCR purification kit and used for yeast cotransformation with the appropriate pCAS plasmid variant. Each 157-bp PCR fragment contained the intron flanked by 50-bp segments homologous to either the small or the large subunit rDNA of yeast, to direct integration by HDR (Fig. 5). The branchpoint and 3’ splice site sequences of the lichen intron match the most common consensus sequences in yeast (Kupfer *et al.* 2004), but the 5’ splice site, GUAAGU, corresponds to a less frequent yeast version. To make the 5’ splice site match the common consensus in yeast, GUAUGU (Kupfer *et al.* 2004), we introduced into the forward primer a one base A>T mismatch with the original lichen sequence (Fig. 5 and Table 1).

**Figure 5.**
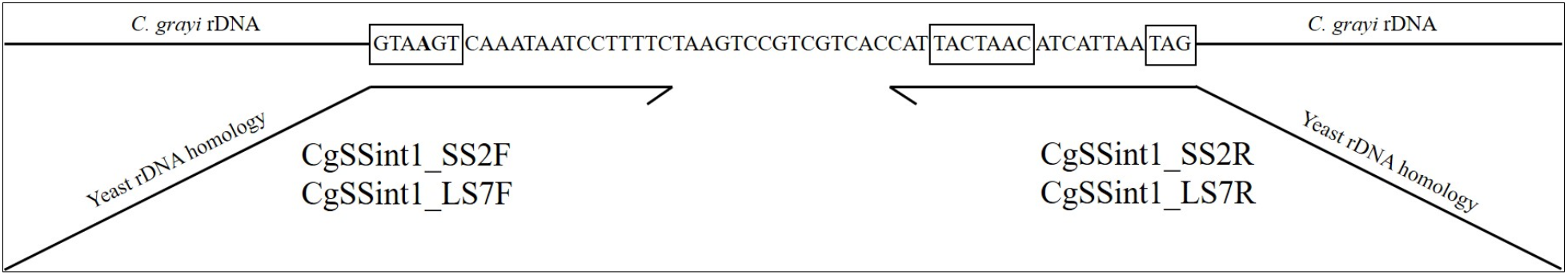
Incorporation of a C. grayi intron into a PCR fragment for integration into yeast rDNA. The top line shows the intron sequence within *C. grayi* rDNA. From left to right, boxes highlight the 5’ splice site, branchpoint, and 3’ splice site consensus sequences, respectively. The bolded A in the 5’ splice site was changed to a T through primer CgSSint1_SS2F or CgSSint1_LS7F. The primer regions annealing to the intron are drawn parallel to the intron, with arrowheads indicating the 3’ ends. Tilted are the 50-bp regions homologous to yeast rDNA, which allow intron integration by HDR into yeast rDNA cut by Cas9 within the homologous regions. Not drawn to scale.

### Yeast transformations, colony PCR, test of plasmid loss

Single intron insertion yeast strains MSS2 and MLS7 were obtained from wildtype strain YJ0 by cotransformation, respectively, with pCAS-SS2 + SSU intron fragment and pCAS-LS7nc + LSU intron fragment. The double insertion strain Md was obtained from strain MSS2 by cotransformation with pCAS-LS7 + LSU intron fragment. We also performed transformations using plasmids pCAS-SS2 or pCAS-LS7nc each by itself, to screen for small mutations around the Cas9 cut sites rather than intron insertions. The protocol by (Ryan *et al.* 2016) was followed for competent cell preparation and transformation. In each cotransformation, 100 μl competent cells were combined with 1 μg pCAS plasmid derivative and 5 μg intron fragment. Transformants were selected on G418 YEPD plates incubated at 30°C. Individual transformants were screened by colony PCR using primers flanking each expected insertion/small mutation site (YrDNA10F/YrDNA18R for the small subunit and Cg28S_F2/Cg28S_R1 for the large subunit, Table 1) and sequencing of PCR products. Thermocycling conditions were as described in Table 2E. After prolonged growth in YEPD lacking G418, yeast transformants were checked for plasmid loss using the same colony PCR protocol with plasmid-specific primers PCR Control F and PCR Control R (Table 1). All yeast mutant strains used in the experiments had lost the Cas9 plasmid.

### Yeast RNA extraction, cDNA preparation

To demonstrate intron splicing, total RNA was extracted from the MSS2 transformant, bearing the intron in the 18S rRNA repeats. A 10 ml overnight culture was pelleted, the pellet was frozen with liquid N_2_ and thoroughly ground in a mortar with liquid N_2_. Total RNA was extracted using the RNAqueous kit (ThermoFisher Scientific). RNA was quantified by NanoDrop (ThermoFisher Scientific) and quality was assessed by agarose gel electrophoresis. Using Superscript III RT (ThermoFisher Scientific), reverse transcription was performed at 50°C for 120 min in a 25 μl volume with 100 ng RNA and the reverse primer YrDNA18R (Table 1), whose 5’ end is 111 bases downstream from the intron insertion site. The cDNA was amplified with primers YrDNA10F and YrDNA18R (Table 1). PCR conditions were as described for yeast colony PCR (Table 2E). The PCR fragment sizes with and without intron are 290 and 233 bp respectively. Correct splicing was confirmed by sequencing.

### Relative rDNA copy number determination

We used qPCR to determine the relative number of rDNA copies in the mutants *vs*. the wildtype strain. Each strain was grown in 10 ml YEPD at 30°C to early stationary phase, and DNA was extracted (Hoffman and Winston 1987). DNA concentration was estimated by Nanodrop (ThermoFisher) and fine-tuned by measuring band intensity on gels. The change in rDNA copy number in MSS2 and MLS7 relative to YJ0 was assessed (Supplementary File 1) with the ΔΔCt method as modified by (Pfaffl 2001) to allow for different amplification efficiencies between reference and target gene. As target we used a section of SSU rDNA, amplified with primers Y-SSUqF2 and Y-SSUqR2, for an amplicon size of 102 bp. As reference we used the single-copy ACT1 gene, amplified with primers Y-Act1qF4 and Y-Act1qR4, for an amplicon size of 87 bp. In each of two replicate qPCR experiments, each DNA was tested in triplicate with the SSU primers and with the ACT1 primers. Standard SybrGreen qPCR reactions were run in 96-well plates in a Bio-Rad Chromo 4 machine, with Opticon Monitor software version 3.1. Cycling conditions are in Table 2F. Amplification efficiencies, 1.55 for SSU and 1.47 for ACT1, were calculated by the Opticon Monitor software.

### Growth Assay

Growth assays were done on YJ0, MSS2, MLS7, and Md. For each strain, a fresh agar plate culture was used to prepare a liquid suspension in YEPD with a concentration between 10^3^-10^5^ cells/ml, assuming an OD_600_ of 1 = 3 × 10^7^ cells/ml. Twelve 150 μl aliquots of the suspension were loaded in three groups (biological replicates) of four technical replicates on two adjacent columns of a 96-round-bottom well plate. To generate growth curves, an automatic plate reader (Tecan) was used to record the OD_600_ at 30°C every 15 minutes over a period of 96 hours. The plate was shaken for 60 seconds before each measurement.

### Desiccation Assay

We modified in two ways the method by (Welch and Koshland 2013). Desiccation was performed in the flow hood rather than in the speedvac and colonies were counted on agar plates rather than directly in the liquid culture wells. All procedures were sterile and media were YEPD. Growth was at 30°C, and liquid cultures were shaken at 255 rpm. Fresh overnight liquid cultures of YJ0, MSS2, MLS7 and Md were diluted to an OD_600_ between 0.1 and 0.25 and grown to 0.5 OD_600_. For each strain, duplicate 1 ml aliquots were pelleted in microfuge tubes. Pellets were resuspended in 1 ml of H_2_O, re-pelleted, and all the water was carefully removed with a pipette. One of the duplicate aliquots (designated as “undesiccated”) was immediately resuspended in 100 μl medium, serially diluted 10x, and three 5 μl replicates from each dilution were spotted on an agar plate. The other duplicate aliquot (“desiccated”) was laid against the sterile air flow in a laminar flow hood to dry for 24 hours, resuspended in 100 μl medium, serially diluted and plated the same way as the undesiccated replicate. The number of survivors in each case was estimated by counting colonies at the highest countable dilutions and extrapolating to the number of viable cells in the starting suspension. Desiccation resistance was defined as the fraction of viable cells in the desiccated *vs.* undesiccated sample. Three to six biological repeats were performed with each strain. P-values were calculated with a two-sided Student’s t-test.

### Microscopy

Budding frequency was measured in a haemocytometer with samples from mid-log phase cultures in YEPD liquid medium. For photography, diluted cell suspensions were plated on YEPD and incubated at 30°C. The YJ0 wildtype was photographed after overnight growth, the mutants after 48-72 hrs. A 1.5 × 1.5 square of agar was cut out of the plate, placed onto a microscope slide, and 10 μl water were placed on the surface followed by a cover slip.

### Intracellular trehalose measurement

The method was essentially as described by (Tapia *et al.* 2015), and (Gibney *et al.* 2015). Each strain was grown at 30°C in YEPD liquid medium to an OD_600_ between 0.3 and 0.5. A volume containing 10^7^ cells was harvested, cells were pelleted, resuspended in 2 ml ice-cold water and re-pelleted. Each pellet was resuspended in 250 μl of 0.25 M sodium carbonate, and the suspension was transferred to a 2-ml screwcap tube and stored a −80°C until trehalose extraction. Trehalose was extracted from cells by incubating the tightly sealed samples at 98°C for 4 hours, with occasional mixing. Samples were stored at −20°C. For trehalase treatment, each 250 μl sample was mixed with 150 μl of 1M acetic acid and 600 μl 0.2 M sodium acetate, cell debris were pelleted, and 250 μl were removed from the top of the sample and stored at −20°C (untreated control). To the remaining 750 μl, 0.35 μl trehalase (Megazyme, at 4.2 units/μl) were added for a final concentration of 2 units/ml, and reactions were incubated overnight at 37°C on a rotating wheel. After pelleting cell debris, the supernatant (750 μl) was transferred to a new tube to be assayed for glucose either immediately or, if convenient, after storage at −20°C, using the Glucose (GO) assay kit from Sigma Aldrich. For each assay, 220 μl of sample were mixed with 440 μl kit reagent in a 15-ml glass test tube, capped and incubated at 37°C for 30 minutes, and 440 μl of 6M sulfuric acid were added in a fume hood with a 1-ml glass pipette. Absorbance at 540 nm was measured in 1-ml plastic cuvettes. The untreated controls were used to measure background glucose, water samples were used as blanks and glucose standards to determine concentration.

## Results

### In the lichen mycobiont, a group I intron is still present in a post-transcriptional rRNA processing intermediate

We assessed by Northern the *in vivo* splicing pattern of intron-3, a group I intron located in the 18S gene of the *C. grayi* lichen fungus (the third from left in Figure 1). Total RNA extracted from the mycobiont grown on MY plates was run on a gel, transferred onto a membrane, and hybridized to an intron-3 specific probe (Fig. 6A). The Northern blot showed two RNAs reacting with the probe, one at ~0.3 Kb and one at ~3Kb. The 0.3 Kb band corresponds to the size of the spliced 310-base intron. The 3 Kb band, significantly larger than the mature 18S rRNA, shows that intron-3 is still present in a post-transcriptional rRNA processing intermediate. The slight smear under the 3 Kb band indicates that it is being partly degraded. The 3 Kb intermediate is likely to be the mycobiont analog of a major yeast 18S rRNA precursor known as “23S” (Klinge and Woolford 2019), which includes the entire 5’ ETS, the 18S sequence, and extends to the A3 cut site in ITS1 (see Figure 3 for a map). Figure 6B schematizes how A3 processing in the lichen fungus would yield the 23S-like intermediate containing intron-3, and how the presence of the intron would interfere with the ongoing assembly either by delaying it through splicing or by removing some of the intermediate by degradation (see Discussion). Figure 6C depicts for comparison the case of a self-splicing fungal rDNA intron which removes itself co-transcriptionally and quickly from nascent rRNA without significantly interfering with ribosome assembly.

**Figure 6.**
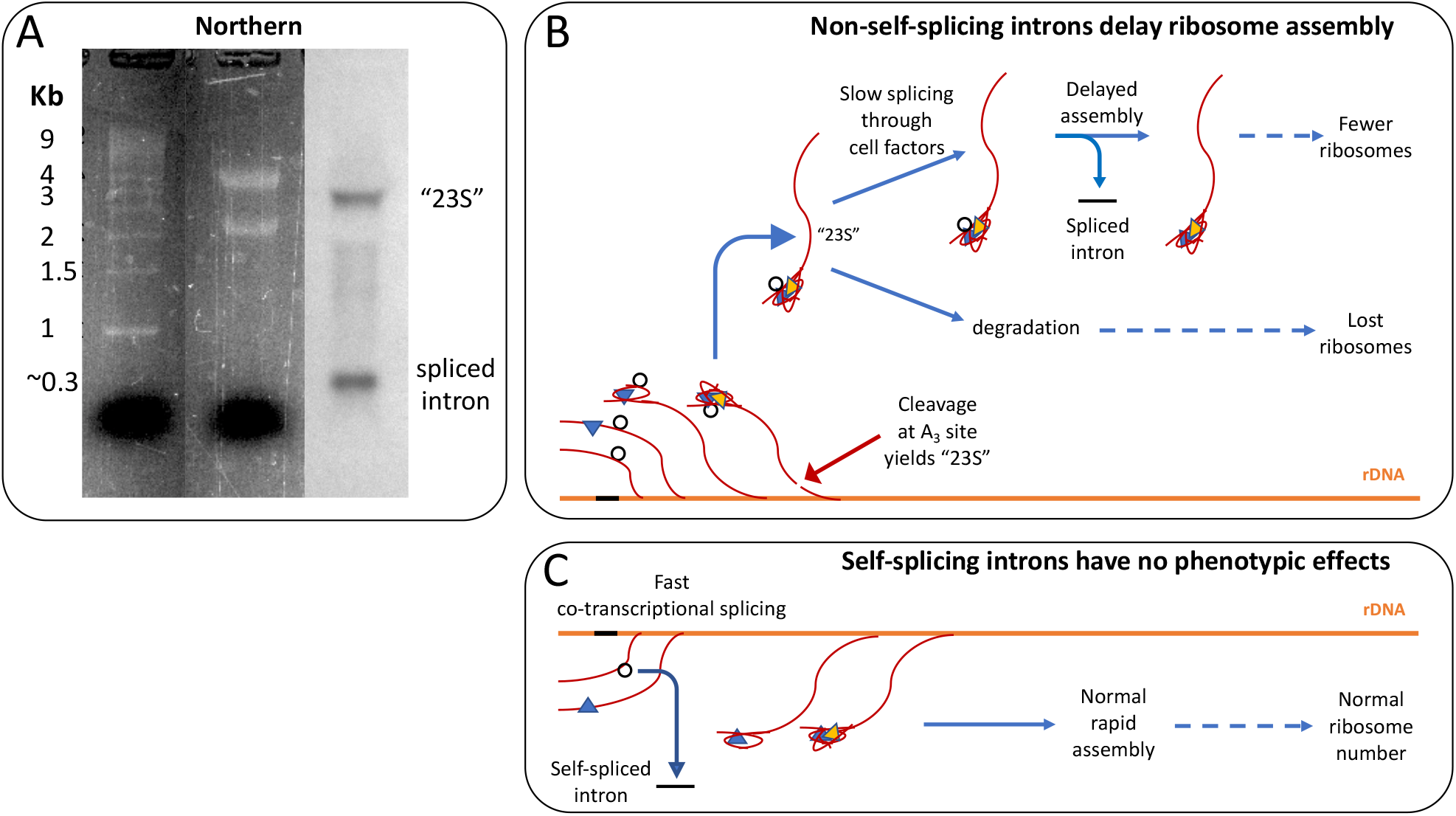
Northern blot and interpretation of the splicing pattern of a degenerate group I rDNA intron (intron-3) in the *C. grayi* lichen fungus. **A.** composite image of the Northern. The first two lanes represent the SybrSafe-stained agarose gel, with size standards in the first and total RNA in the second lane, with prominent 25S and 18S bands. The third lane shows hybridization of the intron-3 probe to the RNA from the second lane transferred onto a membrane. **B.** Schematic of the process by which the 23S-like intermediate is formed during transcription of rRNA (red curvy lines emanating from the orange rDNA). The inserted intron is depicted either as a black segment on rDNA, or as an unspliced black circle on rRNA. Colored triangles schematize proteins assembling with rRNA into pre-ribosomal particles (Fig. 3). **C.** Schematic for comparison of the rapid cotranscriptional removal of a self-splicing intron from nascent rRNA.

### CRISPR is effective in introducing introns or base pair changes into all yeast rDNA repeats

Genetic manipulations of lichen fungi are in their infancy (Wang *et al.* 2020; Liu *et al.* 2021). Therefore, we tested the effects of lichen rDNA introns in the model fungus *S. cerevisiae*, whose rDNA is normally intronless. We transformed yeast with a 57 base-pair spliceosomal intron (first on left in Figure 1) from the rDNA of the lichen fungus *Cladonia grayi* (Armaleo *et al.* 2019). The splicing signals overlap with those of yeast mRNA introns (Fig. 5) and thus are expected to be recognized by yeast spliceosomes. CRISPR-Cas9 technology was seen as the only available way to simultaneously insert the intron into 150 repeats of yeast rDNA (Chiou and Armaleo 2018; Sanchez *et al.* 2019) because of the powerful selection of CRISPR against unmodified rDNA. Our method was developed to insert introns into the 18S and 25S rRNA genes by Homology Directed Repair (HDR), but we list here for the record also the base pair substitutions produced through Non-Homologous End Joining (NHEJ) during this work (Supplementary File 2).

Three intron-bearing mutants were constructed, with the intron stably inserted either in all copies of the 18S (SSU) gene, of the 25S (LSU) gene, or of both. The selected yeast insertion sites corresponded to two *C. grayi* intron sites and had appropriately located PAM sequences (Fig. 7). Intron insertions were obtained by cotransformation of yeast with a Cas9-gRNA plasmid and the corresponding intron containing fragment. Single intron insertion mutants MSS2 and MLS7 contain the intron in the 18S or 25S rRNA gene at positions 534 or 2818 respectively (Fig. 7) The double insertion mutant Md contains the intron at those positions in both genes. All mutants were verified by PCR (Fig. 8) and sequencing; additionally, correct splicing in MSS2 was confirmed by sequencing the mature rRNA. Regardless of whether we inserted introns or produced base-pair mutations around the Cas9 cut sites, the introduced changes appear to involve all repeats as suggested by the absence of heterogeneous peaks at the mutation sites in the sequencing chromatograms (Supplementary File 2) and by the absence of intronless bands in PCRs spanning the intron insertions (Fig. 8). In addition, rDNA introns are stably inherited, which also suggests that no intronless repeats were left to act as recombination seeds and restore the wildtype intronless configuration. Spreading an intron to all yeast rDNA repeats was also achieved by Volker Vogt’s group (Muscarella and Vogt 1993; Lin and Vogt 1998) with a method foreshadowing our use of CRISPR-Cas9 to the same end: to study intron endonucleases, they introduced functional group I rDNA introns from *Physarium polycephalum* or *Tetrahymena thermophila* into yeast rDNA.

**Figure 7.**
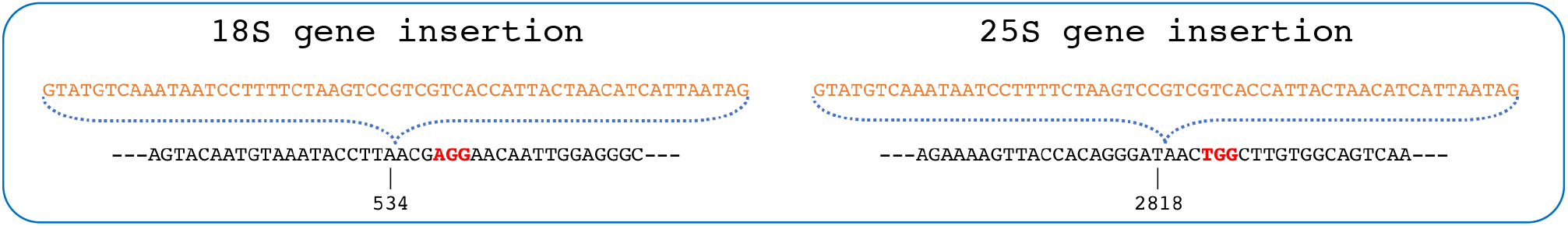
Intron insertion sites in the 18S and 25S rRNA genes of yeast. Only sense-strand sequences are shown, 5’ ends on the left. PAM sequences are marked red. The intron is colored orange and its insertion points by HDR are indicated by the blue-dotted brackets. The vertical dash marks the nucleotide 5’ to the Cas9 cut site on the rDNA sequence and the numbers indicate the corresponding positions in the mature rRNA.

**Figure 8.**
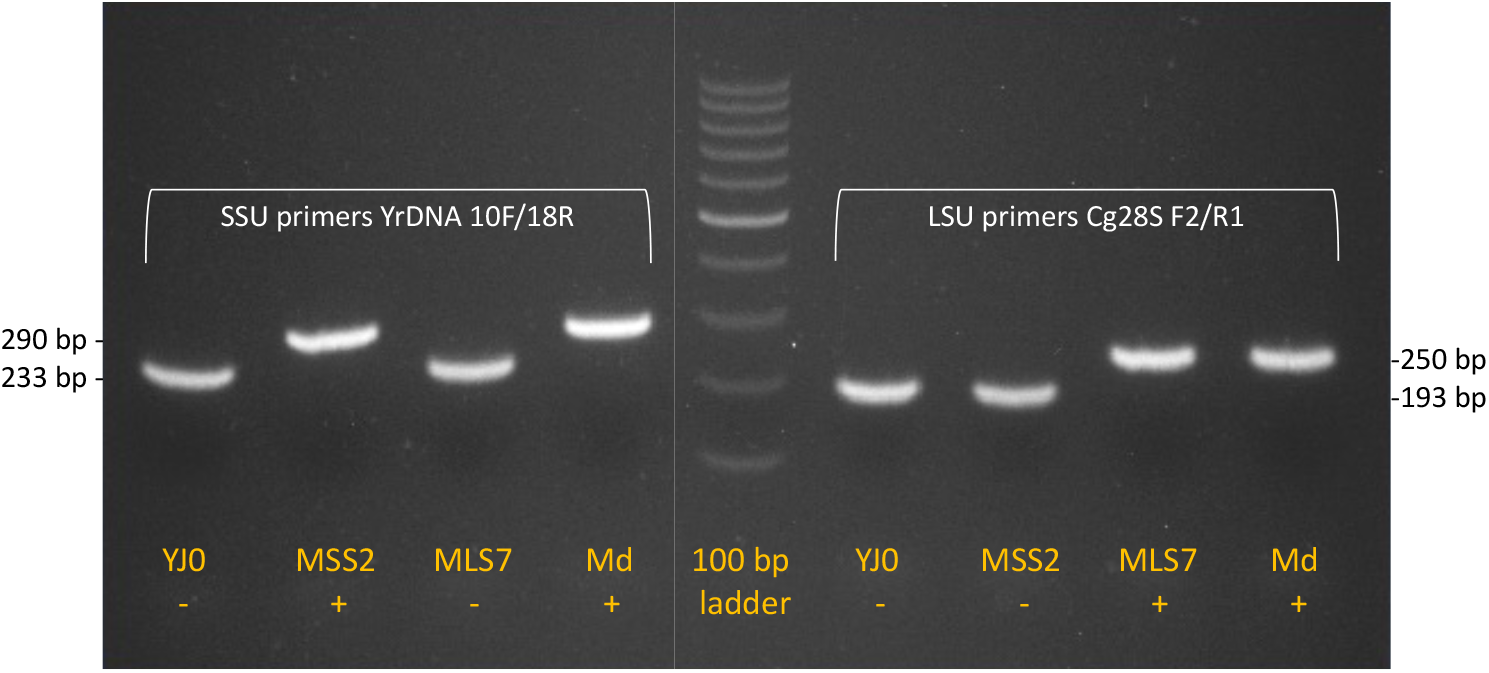
Agarose gel showing intron location in the yeast strains. Primer pairs (white lettering) flanking each intron insertion site were used for PCR with the DNA from the four strains (orange lettering). For each primer pair, the sizes (base pairs) of the fragments with or without the intron are indicated on the side. Intron absence/presence in each fragment is labeled −/+. The sizes indicate that the wildtype strain YJ0 has no introns, MSS2 has the intron only in the SSU gene, MLS7 has the intron only in the LSU gene, Md has the intron in both genes. The lanes with the intron fragments contain no discernible trace of intronless fragments, suggesting that the introns spread through all rDNA repeats.

### Small nucleolar RNAs may interfere with Cas9 when targeting rDNA

Despite two attempts, CRISPR-mediated intron insertion into the rDNA of the YJ0 wildtype using the large subunit guide sequence LS7 (matching the sense strand) failed. It was however successful when we used the LS7nc guide sequence (matching the template strand at the same location). This made us suspect that an endogenous yeast sequence serendipitously complementary to the LS7 guide RNA could have inhibited LS7-Cas9. A small nucleolar RNA (snoRNA) was considered a likely culprit. During yeast rRNA synthesis and ribosome assembly, a number of specific ~100- to 1000-nucleotide long snoRNAs bind as ribonucleoprotein complexes to complementary sequences on the rRNA, leading to post-synthetic nucleotide modifications in those sequences (Dupuis-Sandoval *et al.* 2015). A snoRNA (snR38) with an 11-base complementarity to the 3’ terminal region of the LS7 guide sequence (Fig. 9A) is in fact listed in the yeast snoRNA database (https://people.biochem.umass.edu/fournierlab/snornadb/main.php). Although we do not demonstrate this directly here, the likelihood that snR38 inhibited Cas9 by binding to LS7 is supported a) by the fact that the complementarity is in the snoRNA region that is meant to bind directly to rRNA to methylate it, b) by the successful insertion of the intron at the same position when using the LS7nc guide which is not complementary to snR38 (Fig. 9A and B), and c) by the absence of a snoRNA complementary to the SS2 guide sequence which mediated the successful insertion of the intron into the SS2 site. Figure 9B also shows that the exact insertion point is determined by the placement of the intron within the cotransforming PCR fragment, even if it is a few bases removed from the actual Cas9 cut site.

**Figure 9.**
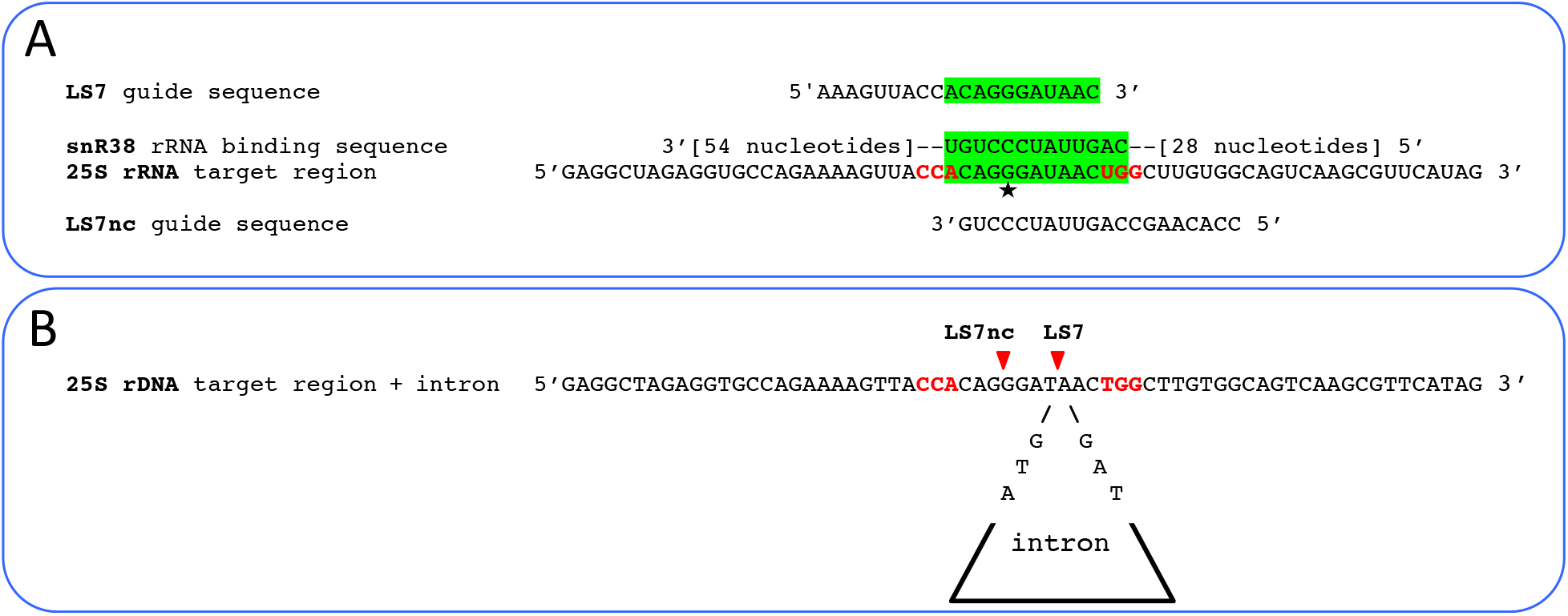
Cause of the likely inhibition of Cas9 by snR38, and intron targeting to the LSU gene. **A.** Complementary binding between the RNA-binding region of snR38 and the LS7 guide RNA is likely to have inhibited the Cas9-LS7 complex. RNA complementary regions are highlighted green; nucleotides corresponding to a PAM sequence on either strand are in red; the star marks rRNA nucleotide 2815, targeted for O-methylation by snR38; Cas9 was not inhibited when used with guide RNA LS7nc, which could not bind to snR38. **B.** The intron was inserted at the LS7 site using LS7nc guide RNA. Red arrowheads indicate the Cas9 cut sites (at positions 2814 for LS7nc and 2818 for LS7) corresponding to the two opposite-strand PAM sites (in red). While LS7nc directed Cas9 to cut the site on the left, homology directed repair integrated the intron in the rDNA in correspondence of the LS7 site at position 2818, where the intron was located in the PCR fragment used in cotransformation.

### Introns decrease rDNA copy number, inhibit growth, and modify cell morphology

To assess whether intron presence affects rDNA copy number, we used the ΔΔCt qPCR method by (Pfaffl 2001) to determine the relative number of rDNA copies in mutants *vs.* wildtype. Copy numbers in each of the single-intron mutants MSS2 and MLS7 decreased to about 53% of the YJ0 wildtype (Supplementary File 1). The qPCR data from the double-intron mutant Md were uninterpretable for unclear reasons. The intron-bearing mutants produced small, slow-growing colonies and their growth rates in liquid culture were 4-6 times slower than that of YJ0 (Fig. 10). It is noteworthy that the single insertion in the LSU region caused a much stronger growth inhibition than the one in the SSU region, pointing to the importance of intron context. The slowdown appears to lengthen the cell cycle in G1, as the strains’ budding frequency is lowered from 65.6% in the wildtype YJ0 to 27.4% (MSS2), 43% (MLS7), and 39.7% (Md). On average, cells of the intron-bearing strains are larger than those of the wildtype. The cell morphology of MLS7 and Md is also frequently abnormal (Fig. 11) and may have inflated the bud counts in these strains.

**Figure 10.**
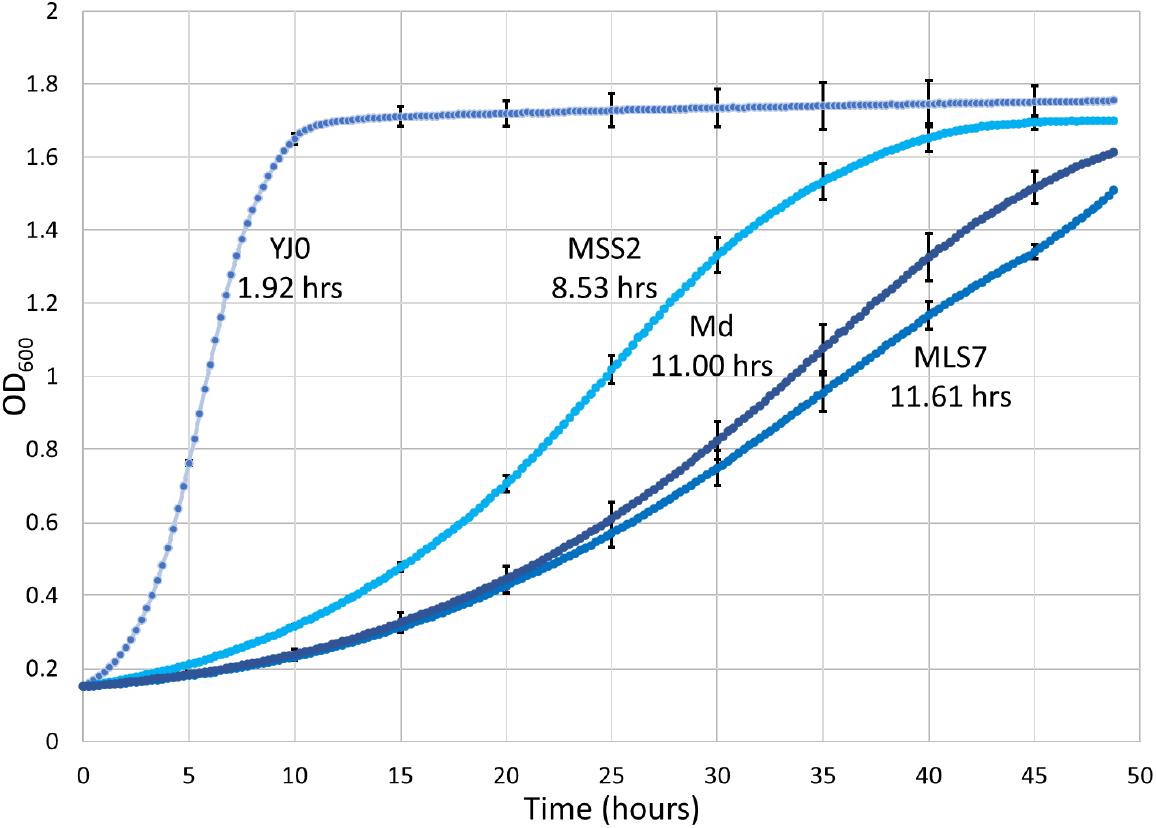
Growth curves for the four strains. Curves represent the means of three biological replicates and four technical replicates for each strain. Bars represent the SD. The numbers next to the curves are the doubling times for each strain.

**Figure 11.**
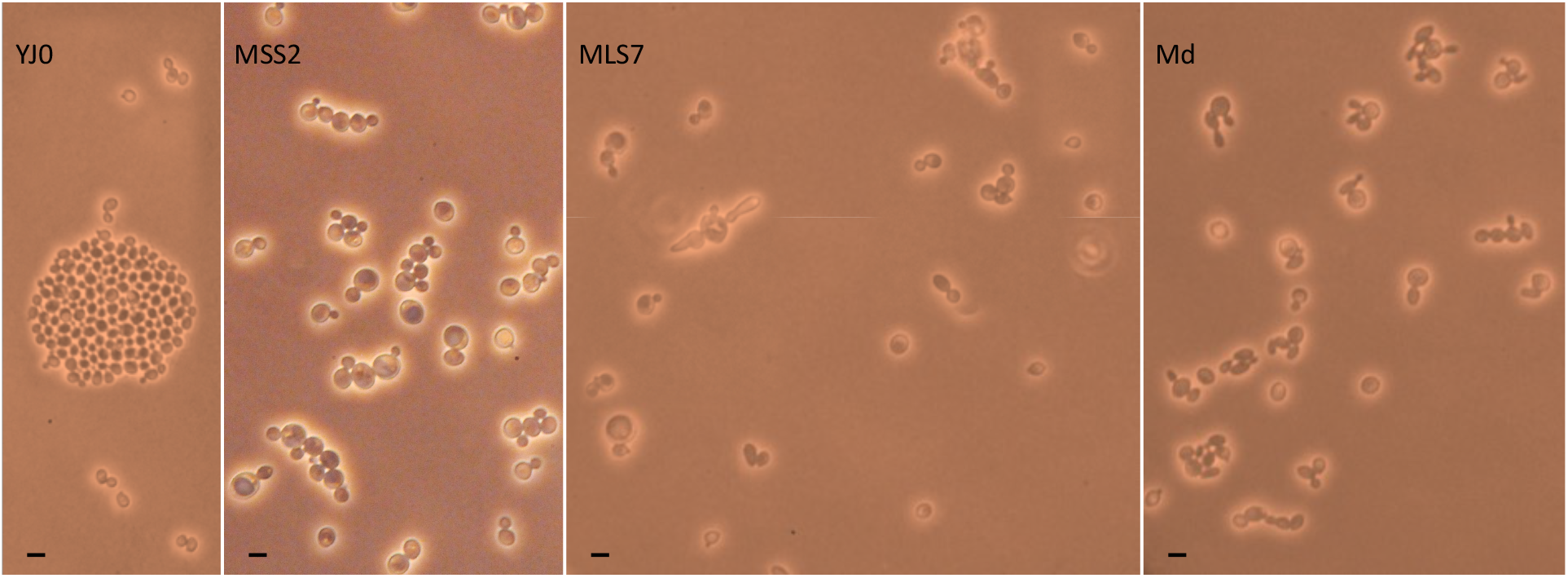
Cell size and morphology. Cells were photographed growing on the surface of freshly seeded agar plates. Bar = 10μ. Intron-bearing cells are on average larger than the YJ0 wildtype cells. Several MLS7 and Md cells have aberrant morphologies. Due to its faster growth, YJ0 quickly forms many incipient colonies, one of which is visible here.

### Introns enhance desiccation tolerance in *S. cerevisiae*

We measured desiccation tolerance in the wildtype and the three intron-bearing mutants using a method modified from (Welch and Koshland 2013). Each strain was grown to mid-log phase, two identical samples were removed, and one was subjected to desiccation while serial dilutions of the other were plated. One day later the desiccated sample was resuspended and serial dilutions were plated. Desiccation tolerance (Fig. 12) for each strain is expressed as the ratio of live cells (scored as colonies) in desiccated *vs.* undesiccated samples of that strain. Figure 12A shows the resistance of individual biological replicates. Three biological replicates were assayed for YJ0 and Md, and five and six biological replicates were assayed for MSS2 and MLS7, respectively. The variation between replicates might be due to small between-sample differences in drying speed and/or to the stochastic behavior of the biochemical networks underlying stress tolerance (see Discussion). Figure 12B shows the same data plotted as averages of the biological replicates. For YJ0, the average absolute survival was 1.5 × 10^−5^. The average absolute (and relative) survival ratios for MSS2, MLS7, and Md were 6.3×10^−4^ (42), 4.5×10^−3^ (300), and 2.6×10^−2^ (1700). The position and number of introns affects desiccation resistance, paralleling their effect on growth (Fig. 10). MLS7 with the intron in the large subunit is more desiccation resistant than MSS2 with the intron in the small subunit. Md, the strain with introns in both subunits, is the most resistant.

**Figure 12.**
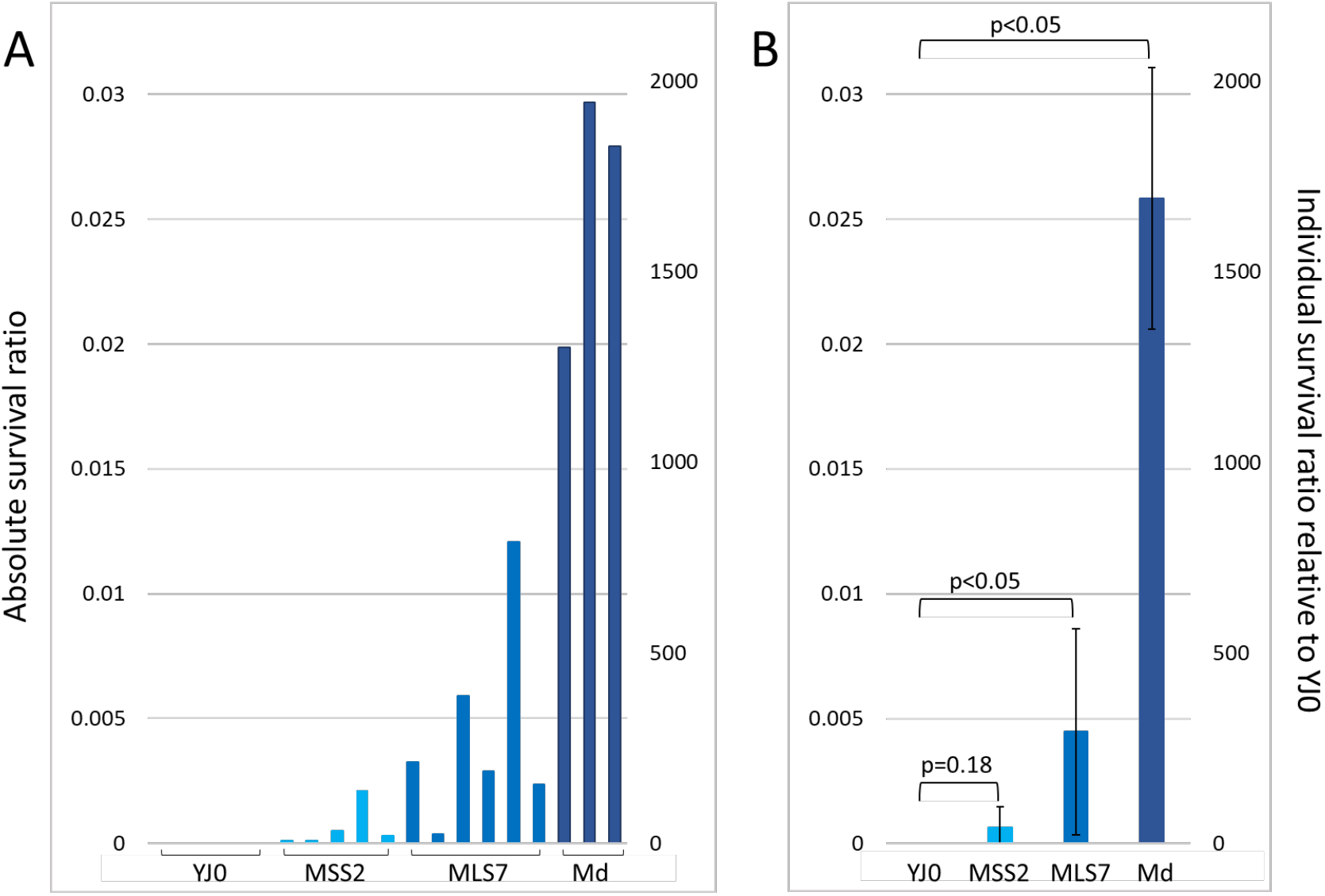
Desiccation resistance of the four strains. The Y axes are the same in both panels. Absolute survival ratios represent the fraction of cells surviving desiccation within each strain. Relative survival ratios represent each strain’s resistance relative to that of the YJ0 wildtype. **A.** Resistances of individual biological replicates. **B.** Same data averaged over biological replicates; SD in black; brackets indicate p values for mutant *vs*. wildtype ratios.

### Introns induce trehalose biosynthesis in the double mutant

In some organisms, including yeast, intracellular concentrations of the disaccharide trehalose are positively correlated with desiccation tolerance (Koshland and Tapia 2019). We therefore measured whether intracellular trehalose accumulates in our mutants during normal growth. The procedure involves extraction of trehalose from a fixed number of mid-log phase cells grown in YEPD medium, trehalase treatment to split trehalose into its two glucose constituents, and a colorimetric assay that measures glucose concentration. The results (Fig. 13) indicate that trehalose was undetectable not only in the YJ0 wildtype, but also in the single-intron mutants MSS2 and MLS7, despite their increased desiccation tolerances. Trehalose increased to detectable levels only in the Md strain. Md carries two introns, one per subunit in each rDNA gene copy, and displays the highest desiccation tolerance (Fig. 12).

**Figure 13.**
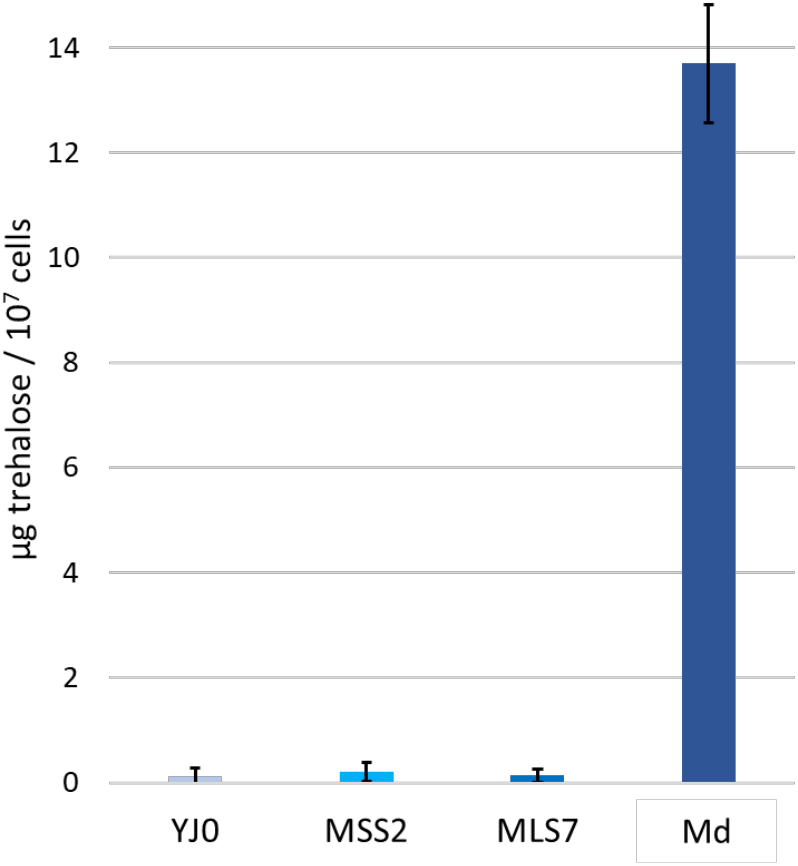
Intracellular trehalose in the four strains. Amounts are expressed /10^7^ cells /ml of assay. For each strain, the SD is calculated on 9 samples (three biological replicates, each with three technical replicates).

## Discussion

The successful use of CRISPR to mutagenize the entire array of rDNA repeats in yeast has been described and discussed (Chiou and Armaleo 2018; Sanchez *et al.* 2019). The interesting, albeit peripheral issue of snoRNA interference with Cas9 described in Results is not further discussed here. In our first experiment, we used the *C. grayi* lichen fungus to assess, using a Northern blot, the splicing pattern of a degenerate group I intron naturally present in its rDNA. All other experiments were done by approximating a lichen-like nuclear rDNA intron arrangement in yeast. Using CRISPR, a lichen spliceosomal intron was introduced at one or two positions into each yeast rDNA repeat (Figs. 7 and 8). We could not have used a lichen degenerate group I intron, as yeast does not have the machinery to splice it out. Bending the general rule that spliceosomes operate only on Polymerase II transcripts, yeast successfully removed the lichen spliceosomal introns from the Polymerase I rRNA transcripts, allowing us to model in yeast the effects of lichen rDNA introns. The goal was to test in yeast whether the inability of lichen rDNA introns to self-splice may lead to slow growth and desiccation tolerance.

### In the lichen mycobiont, delayed group I intron splicing is likely to slow ribosome assembly

The ~300-base size of the smaller band in the Northern blot (Fig. 6A) and the absence of a signal from the mature 18S rRNA indicate that the degenerate group I intron-3 in the rRNA of the *C. grayi* mycobiont is correctly spliced out as a complete ~300-nucleotide fragment. However, the ~ 3 Kb size of the larger band indicates that the splicing is delayed enough for the intron to persist within an accumulating precursor of the 18S rRNA. The size of this intron-bearing precursor is slightly larger than that of a 2.8-Kb yeast rRNA intermediate (23S) observed when yeast grows slowly due to mutational or nutritional stress (Venema and Tollervey 1999; Talkish *et al.* 2016; Klinge and Woolford 2019) and is an indication that under those circumstances rRNA processing is delayed until after transcription of the entire 35S gene product is terminated (Talkish *et al.* 2016). The yeast 23S precursor contains a fully unprocessed 5’ ETS, the 18S sequence, and ends at the A_3_ processing site in ITS1 (see Fig. 3 for a map). The 23S precursor is part of the SSU processome and its 5’ETS region needs to be rapidly removed for ribosome assembly to continue (Klinge and Woolford 2019). Therefore, accumulation of an unprocessed intron-bearing 23S-like precursor in the mycobiont supports our hypothesis that intron presence delays ribosome assembly in lichens (Fig. 6B). The Northern shows also that the precursor is partly degraded (Fig. 6A). The 23S rRNA precursor accumulating in yeast under slow-growth conditions is frequently degraded and associated with decreased levels of mature 18S rRNA, *i.e.* with ribosome depletion (Venema and Tollervey 1999; Talkish *et al.* 2016). Degradation of stalled rRNA processing intermediates in yeast due to an rDNA intron defective in self-splicing has also been reported for the large subunit by (Jackson *et al.* 2006). The processing pattern in fast-growing yeast (conserved in other fast-growing fungi) is quite different, as most processing of the 18S rRNA precursor region occurs through rapid co-transcriptional removal of the entire 5’ETS and most of ITS1 (Fernández-Pevida *et al.* 2015; Klinge and Woolford 2019), with no accumulation of 23S rRNA (Fig. 6C). In conclusion, accumulation of an intron-3 bearing 23S-like intermediate and its partial degradation in the *C. grayi* fungus strongly imply that intron-3 and most other lichen rDNA introns delay ribosome assembly and reduce ribosome number.

### The slow growth of the yeast mutants suggests that non-self-splicing rDNA introns inhibit ribosome biogenesis constitutively

The large growth decreases in the intron-bearing yeast mutants (Fig. 10) also support our main hypothesis that interference by spliceosomes with the highly cooperative and perturbation sensitive ribosome assembly (Osheim *et al.* 2004) delays rRNA processing and slows growth. The growth decreases depend on position and number of introns, but not in a linear fashion (Fig. 10). The structures of rRNA folding intermediates (Klinge and Woolford 2019), indicate that the 18S gene intron insertion site is located at the surface of the SSU processome, whereas the 25S insertion site is buried deep in the folding LSU pre-ribosomal particle, near the developing entrance to the polypeptide exit tunnel. The interference of intron splicing with LSU assembly might therefore be stronger than with SSU assembly, suggesting an explanation for the growth differential between the two single insertions and also for the similar growth rates of the double insertion and the single 25S rRNA insertion. The growth defects are permanent, as expected for a process co-occurring with rRNA transcription. Also, other phenotypes we observe in the mutants, like rDNA repeat number reduction, G1 delays, and aberrant cell morphologies (Fig. 11), are likely consequences of intron interference with ribosomal assembly. These phenotypes could all result from DNA replication stress (Tripathi *et al.* 2011; Salim *et al.* 2017) perhaps triggered by the interference of intron splicing with the functions of early 60S assembly factors Noc3p and Rix, which are also part of the DNA pre-replicative complex (Zhang *et al.* 2002; Dez and Tollervey 2004; Huo *et al.* 2012).

Besides interfering with rRNA processing through their splicing machinery, rRNA spliceosomal introns may also inhibit growth by competing with their mRNA counterparts for limiting spliceosomal components, directly inhibiting splicing of ribosomal protein (RP) mRNAs and RP synthesis. Competition between the highly expressed intron-rich RP genes, which use the most splicing factors (Ares *et al.* 1999), and other intron-containing genes has been demonstrated in yeast (Munding *et al.* 2013). Even with a reduced number of rDNA genes (70 to 80), the single intron mutants would have roughly 25% more spliceosomal introns in their genome than the wildtype strain. The inhibition of RP synthesis might be deepened if these introns persist as undegraded and linearized RNAs after splicing (Morgan *et al.* 2019; Parenteau *et al.* 2019).

### Induction of desiccation tolerance is stochastic and independent of the onset of desiccation

In yeast, ribosomal synthesis during unstressed growth is enabled by the activity of the TOR and PKA signal transduction pathways. Environmental stress signals repress TOR/PKA, which regulate the two arms of the Environmental Stress Response (ESR) (Gasch *et al.* 2000). One arm involves ~600 genes which are turned down under stress and include ribosomal biogenesis (RiBi) genes needed for rRNA, RP, and assembly factor production. The other involves ~300 genes which are upregulated by stress and are responsible for specific induced defenses (iESR) like the disaccharide trehalose, chaperone proteins and redox effectors.

Extreme desiccation stress quickly deprives cells of water, the most basic ingredient for any life-sustaining process. This makes it critical for desiccation-defenses to be in place before the onset of desiccation. Most cells in an unstressed yeast population will not express the ESR and will die when desiccated. However, due to cell-to-cell stochasticity of transcriptional, translational and post-translational network regulation (Gasch *et al.* 2017), a few cells will express the ESR during unstressed growth in liquid media and survive desiccation. This “bet-hedging” (Levy *et al.* 2012) allows a single-celled, fast-growing organism like yeast to thrive evolutionarily, as variable subsets of the population survive under changing stresses, even if most of the population dies. The average desiccation survival of normal intronless *S. cerevisiae* growing in rich liquid media is ~10^−6^. That means that in the bet-hedging process only one in a million cells has serendipitously activated, before desiccation, the ESR defenses needed to survive. In other yeast experiments that uncovered processes or molecules protecting cells from desiccation damage (Gadd *et al.* 1987; Calahan *et al.* 2011; Welch *et al.* 2013; Tapia *et al.* 2015; Kim *et al.* 2018), desiccation tolerance could be experimentally increased up to 10^6^ fold relative to normal “unprepared” yeast as increasingly larger fractions of cells expressed their defenses before desiccation was applied.

The intron-bearing mutants are 40 to 1700 times more resistant to desiccation than the wildtype strain, depending on intron position and number (Fig. 12). The highest resistance corresponds to only 2.6% of cells surviving desiccation, with bet-hedging still occurring, although resistance is spread to many more cells than in the YJ0 wildtype. Our desiccation results parallel those by (Welch *et al.* 2013), who inhibited ribosomal biogenesis through the TOR pathway in two specific ways, by treating yeast with rapamycin or by deleting the TOR effector SFP1 which is a positive regulator of ribosomal protein and assembly genes. Either method increased desiccation resistance. In addition, they tested 41 temperature sensitive (Ts) ribosome assembly protein mutants which, like our intron mutants, impaired ribosome assembly independently of TOR/PKA. Also like ours, the Ts mutations were expressed in liquid media before desiccation, and all produced various degrees of desiccation resistance, the highest matching the 2-3% resistance of our double mutant.

### Ribosome number reduction and trehalose increases represent two complementary defenses against desiccation, the first constitutive and the second inducible

While the slow growth of the yeast mutants is likely a direct, even if not linear, consequence of intron-mediated RiBi repression, the connection between the latter and increased desiccation tolerance is less clear. However, viewing the trehalose results within the context of rDNA introns sheds some light on the issue. The disaccharide trehalose is a primary anti-desiccation molecule in several anhydrobiotes (Koshland and Tapia 2019). In yeast it is induced through the iESR with several other defenses and reduces protein misfolding and aggregation *in vitro* and *in vivo* (Kim et al. 2018). Trehalose was assayed in the intron-bearing mutants as they grew slowly in rich liquid media in absence of environmental stress.

Intracellular trehalose concentration, naturally or experimentally induced, is generally proportional to desiccation resistance in intronless yeast (Gadd *et al.* 1987; Tapia and Koshland 2014). In our strains, despite their increased desiccation tolerances, trehalose remained undetectable in both single-intron mutants, and was detected only in the double-intron mutant Md at 14 μg/ml (Fig. 13). This concentration would be too low in normal, intronless yeast to account for the 2.6% desiccation tolerance shown by Md (Fig. 12). Yeast tolerances around 1% require trehalose concentrations between 150 μg/ml (Fig 2 in (Tapia *et al.* 2015)) and 700 μg/ml (Fig 2 in (Tapia and Koshland 2014)), depending on the experiment. Our trehalose results resemble findings by (Calahan *et al.* 2011) who showed that desiccation resistance remained high in slow-growing yeast after diauxic shift even without trehalose (eliminated through pathway mutations).

This indicates that, whether yeast grows slowly because of rDNA introns or because of depletion of fermentable carbon sources, factors other than inducible defenses like trehalose can elicit desiccation tolerance. We believe that data in the literature strongly suggest that a central component of those “other factors” is the constitutive reduction of cytoplasmic ribosome number. (Welch *et al.* 2013) showed that TOR dependent and independent inhibition of ribosome function increases desiccation tolerance; (Pestov and Shcherbik 2012) showed through rRNA quantitation that rapamycin inhibition of TOR during yeast exponential growth degrades ~50% of ribosomes; (Delarue *et al.* 2018) demonstrated directly that inhibition of the TOR pathway by rapamycin increases cytoplasmic diffusion rates by decreasing ribosome numbers by 40-50%. They also showed that diffusion rates increase without rapamycin while yeast cells enter stationary phase, again indicating a decrease in ribosome number during this transition, shown by (Calahan *et al.* 2011) to produce large increases in desiccation tolerance. Ribosome depletion during entry into stationary phase was also observed directly at the rRNA level by (Talkish *et al.* 2016). We therefore hypothesize that permanent RiBi repression by rDNA introns produces desiccation tolerance by maintaining a less crowded cytoplasm with fewer ribosomes and proteins. This would reduce protein aggregation, misfolding, and phase separation (Delarue *et al.* 2018), and therefore the load on heat shock proteins and damage to the proteome upon water loss and cell volume shrinkage. Synergy between this constitutive ribosome-based defense and inducible defenses like trehalose, Hsp12 (Kim *et al.* 2018), glycerol and polyols (Dupont *et al.* 2014) might allow lower concentrations of inducible defenses to yield the same protective effects that higher concentrations yield in a normally more crowded cytoplasm. In our yeast mutants, both types of defenses were limited by bet hedging, thus protecting only a subset of cells. Stochastic cell-to-cell variation in ribosome numbers and induced defenses, differentiating desiccation survival between individual cells, is not in contradiction with the constitutive expression of ribosomal introns slowing overall growth of the mutant cell population, which is averaged through time and across all cells.

### Extrapolations on the functions and evolution of rDNA introns in lichens

Unlike the fast-growing, single-celled yeast, lichens are slow-growing, differentiated multicellular networks of hyphae interwoven with consortia of different organisms. Their adaptation to extreme and repeated moisture oscillations must ensure survival of all or most component cells, not just of small subsets as is the case with bet hedging in yeast. We view the acquisition of large numbers of non-self-splicing rDNA introns as a powerful constitutive defense suppressing bet hedging across the entire lichen thallus. Such a constitutive defense based on ribosome reduction might also reduce the amounts of inducible defenses needed, like trehalose, polyols, and Hsps. As the introduction of just two non-self-splicing rDNA introns is quite disruptive to yeast, we imagine that the evolutionary insertion of many introns into mycobiont rDNA was gradual, providing time for selection to turn partial disruptions of RiBi and DNA replication into adaptations. The importance of constitutive stress-defenses in lichens has been also recognized by (Gasulla *et al.* 2021).

How could the degeneracy of lichen Group I introns have evolved? Despite lacking sequences needed for self-splicing, lichen group I introns appear still able to fold into tertiary structures resembling the original ones (DePriest and Been 1992; Bhattacharya *et al.* 2000; Bhattacharya *et al.* 2002; Haugen *et al.* 2004b). It is therefore possible that, while progressively removing catalytically important sections from the introns, natural selection transferred splicing capability to the maturases that originally helped those structures fold. Cloning lichen maturases could verify such a scenario. Why are group I and spliceosomal introns maintained together in lichen rDNA? We think because their primary functions are different: the numerous group I introns primarily interfere with rRNA processing, whereas the spliceosomal introns’ direct interference with rRNA processing is secondary in lichens due to their relatively small numbers; their primary function is to repress ribosomal protein (RP) synthesis by competing for limiting splicing factors with the intron-rich RP mRNAs.

In conclusion, the splicing pattern of a degenerate group I intron in a lichen fungus implies that lichen rDNA introns inhibit ribosome assembly. With yeast, we demonstrate that introduction into its rDNA of a lichen intron requiring spliceosomes for removal strongly enhances desiccation tolerance, also through the likely inhibition of ribosome assembly. We hypothesize that the numerous rDNA introns present in lichen fungi protect the entire mycelium in the thallus essentially by lowering ribosome content and holding the cytoplasm in a permanent state of “molecular frugality” which is less susceptible to desiccation damage. The evolution in lichens of a specific splicing machinery for the degenerate group I introns turned them into repressors of rRNA processing and ribosome assembly, transforming genetic parasites into a new tool against environmental stress; the ectopic rDNA location of spliceosomal introns turned them into repressors of ribosomal protein synthesis to the same end. We think that evolutionarily, the spreading of special introns into the rDNA of lichen fungi was a watershed moment that helped lichens adapt to their permanent exposure to ever-changing environments. The price they paid was slow growth.

## Acknowledgments

We thank John Mercer for useful discussions; Vivian Miao, Paul Manos, John Woolford, Austen Ganley for commenting on the manuscript; Susan May for assistance with the Northern. Duke University Undergraduate Research Support for funding L.C.; Duke University Trinity College of Arts and Sciences for teaching lab funds for D.A.; 73 people, through experiment.com, for research support for D.A. and L.C. The authors have no competing financial interests.

