## Additional File 1 for "Modeling in yeast how rDNA introns slow growth and increase desiccation tolerance in lichens"

### qPCR data elaboration

| Efficiency from both repeats and all samples | |
|---|---|
| SSU primers | Act primers |
| 1,54 | 1,45 |
| 1,58 | 1,49 |
| 1,56 | 1,49 |
| 1,55 | 1,4 |
| 1,54 | 1,42 |
| 1,53 | 1,44 |
| 1,55 | 1,42 |
| 1,53 | 1,45 |
| 1,52 | 1,45 |
| 1,55 | 1,49 |
| 1,57 | 1,49 |
| 1,58 | 1,51 |
| 1,55 | 1,47 |
| 1,57 | 1,47 |
| 1,54 | 1,48 |
| 1,59 | 1,51 |
| 1,59 | 1,52 |
| 1,55 | 1,49 |
| **1,555** | **1,468888889** |
| **Avg E ssu** | **Avg E act** |
| 0,02093407 | 0,034108631 |
| SDV ssu | SDV act |

| | Strain | Ct SSU | Avg Ct SSU | Ct ACT | Avg Ct ACT | Delta SSU YJ0-SS2 | Delta ACT YJ0-SS2 | E_ssu^Dssu | E_act^Dact | Ratio SS2/YJ0 | Delta SSU YJ0-LS7 | Delta ACT YJ0-LS7 | E_ssu^Dssu | E_act^Dact | Ratio LS7/YJ0 |
|---|---|---|---|---|---|---|---|---|---|---|---|---|---|---|---|
| Repeat one | | 17,85 | | 29,73 | | | | | | | | | | | |
| | **YJ0** | 17,96 | 17,70333333 | 29,38 | 29,48 | -4,216666667 | -3,276666667 | 0,157555901 | 0,282980733 | **0,556772538** | -3,89 | -2,89 | 0,181806621 | 0,328437625 | **0,553549919** |
| | | 17,3 | | 29,33 | | | | | | | | | | | |
| | | 21,81 | | 32,82 | | | | | | | | | | | |
| | **SS2** | 21,9 | 21,92 | 32,72 | 32,75666667 | | | | | | | | | | |
| | | 22,05 | | 32,73 | | | | | | | | | | | |
| | | 21,36 | | 32,27 | | | | | | | | | | | |
| | **LS7** | 21,66 | 21,59333333 | 32,58 | 32,37 | | | | | | | | | | |
| | | 21,76 | | 32,26 | | | | | | | | | | | |
| Repeat two | | 17,47 | | 29,2 | | | | | | | | | | | |
| | **YJ0** | 17,63 | 17,26666667 | 29,05 | 29,07 | -3,566666667 | -2,143333333 | 0,20948372 | 0,437908302 | **0,478373482** | -2,116666667 | -0,686666667 | 0,395486113 | 0,767553689 | **0,515255309** |
| | | 16,7 | | 28,96 | | | | | | | | | | | |
| | | 20,49 | | 31,41 | | | | | | | | | | | |
| | **SS2** | 20,86 | 20,83333333 | 31,15 | 31,21333333 | | | | | | | | | | |
| | | 21,15 | | 31,08 | | | | | | | | | | | |
| | | 19,18 | | 29,81 | | | | | | | | | | | |
| | **LS7** | 19,5 | 19,38333333 | 29,96 | 29,75666667 | | | | | | | | | | |
| | | 19,47 | | 29,5 | | | | | | | | | | | |

| | Mutant/YJ0 | Ratios | SS2 AVG | LS7 AVG | Global AVG | SDV | AVG - SDV | AVG + SDV |
|---|---|---|---|---|---|---|---|---|
| repeat one | **SS2/YJ0** | 0,556772538 | 0,51757301 | 0,534402614 | 0,525987812 | 0,0369219 | 0,489065912 | 0,562909712 |
| | **LS7/YJ0** | 0,553549919 | | | | | | |
| repeat two | **SS2/YJ0** | 0,478373482 | | | | | | |
| | **LS7/YJ0** | 0,515255309 | | | | | | |

Based on: Nucleic Acids Res. 2001 May 1; 29(9): e45. doi: 10.1093/nar/29.9.e45

$$\text{Mutant/YJ0 rDNA copies ratio} = \frac{1.55^{(\text{YJ0 SSU ct} - \text{Mutant SSU ct})}}{1.47^{(\text{YJ0 ACT ct} - \text{Mutant ACT ct})}}$$
