## Additional File 2 for "Modeling in yeast how rDNA introns slow growth and increase desiccation tolerance in lichens"

**Point mutations generated though CRISPR are homogenized throughout the rDNA**

Individual base-pair mutations generated by NHEJ were obtained by transforming strain YJ0 only with a pCAS plasmid engineered to target either position 534 in the 18S or position 2818 in the 25S gene (Fig. 1). Several clones had the same mutation and the total number of mutant base configurations we detected was small (four around 18S position 534 and one around 25S position 2818), not surprising since each surviving mutation needs to produce a functional, even if suboptimal, rRNA. Most base-pair change mutants grew more slowly than the parent. Overall, almost all transformants fulfilled the expectation that any repeat with a wild type sequence around the target site would not survive Cas9 attack. However, some transformants had no change around the target site, and actually grew at wild type rates or faster. We did not analyze those further.

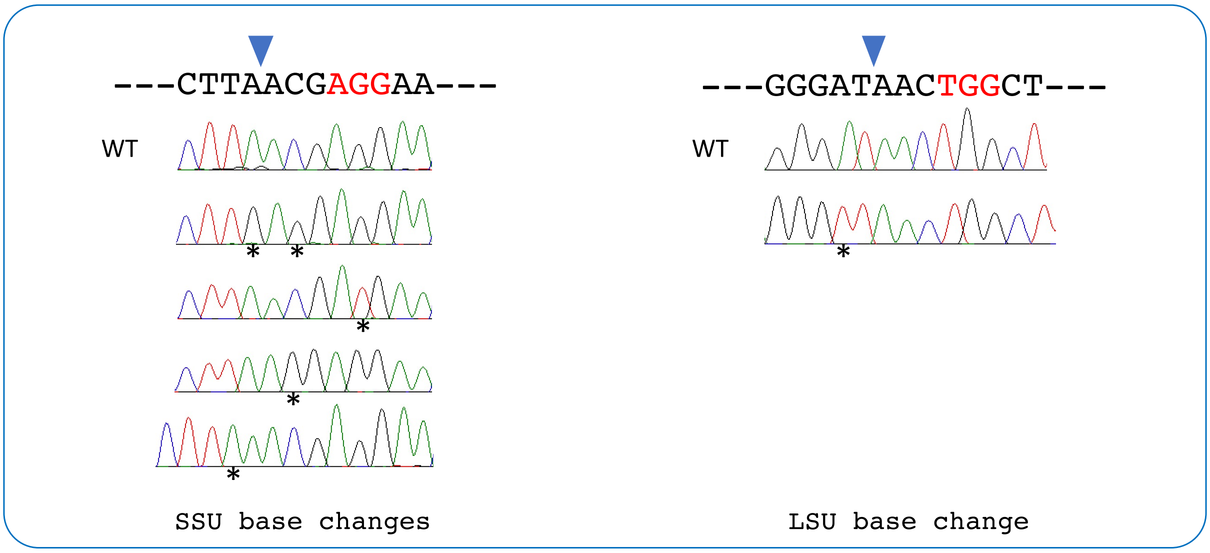


**Figure 1.** Individual base pair changes around the Cas9 cut sites. Only sense-strand sequences are shown, 5' ends on the left. PAM sequences are marked red. Arrowheads point to Cas9 cut sites. The wild type sequence at each site is indicated above the chromatograms of representative yeast transformants bearing base changes (asterisks) produced through NHEJ. The traces show no peak heterogeneity among the sequenced PCR fragments at the mutated sites, suggesting that the same base change spread to all rDNA repeats.
